# Using genome scans to identify genes used repeatedly for adaptation

**DOI:** 10.1101/2022.03.24.485690

**Authors:** Tom R. Booker, Sam Yeaman, Michael C. Whitlock

## Abstract

Adaptation occurring in similar genes or genomic regions in distinct lineages provides evolutionary biologists with a glimpse at the fundamental opportunities for and constraints to diversification. With the widespread availability of high throughput sequencing technologies and the development of population genetic methods to identify the genetic basis of adaptation, studies have begun to compare the evidence for adaptation at the molecular level among distinct lineages. However, methods to study repeated adaptation are often oriented towards genome-wide testing to identify a set of genes with signatures of repeated use, rather than evaluating the significance at the level of an individual gene. In this study, we propose *PicMin*, a novel statistical method derived from the theory of order statistics that can test for repeated molecular evolution to estimate significance at the level of an individual gene, using the results of genome scans. This method is generalizable to any number of lineages and indeed, statistical power to detect repeated adaptation increases with the number of lineages that have signals of repeated adaptation of a given gene in multiple lineages. An implementation of the method written for R can be downloaded from https://github.com/TBooker/PicMin.

## Introduction

Repeated adaptation is ubiquitous across the tree of life. For example, many protein-coding genes are shared among deeply diverged species (e.g. 16S ribosomal RNA genes) with purifying selection acting to maintain the integrity of such genes in distinct lineages. This broadly defined process of repeated adaptation is widespread across many species’ genomes. However, one can also think of repeated adaptation in a narrower sense: when distinct lineages adapt to similar yet novel selection pressures via changes in orthologous genes. As repeatability is the cornerstone of statistical power, observing a gene’s involvement in multiple independent bouts of adaptation can provide stronger evidence that it is truly causal and not just a spurious pattern due to genetic drift. Furthermore, repeated use of the same genes – and repeated lack of use of other genes – in independent bouts of evolution can give insights into the nature of constraints and opportunities in the genotype-phenotype-fitness map (Yeaman et al. 2018) or how the interplay between standing variation, *de novo* mutation, and migration shapes evolutionary outcomes (Lee and Coop 2017).

Various forms of repeated adaptation are increasingly discussed in the population genomics literature, where the same genes or genomic regions are implicated in bouts of adaptation in distinct lineages (e.g. Yeaman et al. 2016; Rennison et al. 2019; Bohutínská et al. 2021; Tittes et al. 2021; Rennison and Peichel 2022). However, it is important to note that distinct selection regimes can give rise to a pattern of repeated adaptation. For example, if multiple species inhabited a given latitudinal temperature gradient, both species could evolve increased cold tolerance through changes in the same gene. Alternatively, one species may evolve reduced cold tolerance while another may evolve increased cold tolerance, but through changes in the same gene. In these two examples, there was a shared pattern of adaptation at the individual genes, although phenotypic evolution went in opposite directions. Referring to such patterns as repeated adaptation we intentionally remain agnostic to the direction of phenotypic change (i.e. divergent versus convergent evolution) or the source of genetic variation (i.e. convergent versus parallel evolution), but the reader should note that terminology varies in the literature (e.g. Lee and Coop 2019).

Depending on the temporal and spatial scale of adaptation, many different summary statistics may be used to quantify a gene’s adaptation within a given lineage, such as *F*_*ST*_, nucleotide diversity (*π*) or genotype-environment associations calculated for each gene (see reviews by Casillas and Barbadilla (2017) and Hoban et al. (2016)). However, for many population genetic summary statistics, it is exceedingly difficult to adequately model the null distribution and calculate accurate *p*-values. This is because the null distributions for population genetic summary statistics may be sensitive to the details of population structure and sample design (Lotterhos and Whitlock 2015) and factors that are idiosyncratic to individual lineages such as recombination rate variation (Booker et al. 2020) and demographic history (Lotterhos and Whitlock 2015; Johri et al. 2020).

To detect repeated adaptation, the results of genome scans in individual lineages are sometimes compared to an arbitrary significance threshold to classify genes or genomic regions as being adapted/non-adapted. With lists of adaptation candidates for each lineage, the overlap among these lists is tested to determine whether it exceeds some null expectation. Such overlap methods have revealed some interesting evolutionary patterns in recent studies; for example, Bohutínská et al. (2021) showed that the extent of repeated adaptation was related to genetic distance among lineages, and Tittes et al. (2021) showed that repeated adaptation between maize and teosinte is often facilitated by the migration of alleles across populations. While these methods yield insights into genome-wide patterns of repeated adaptation, they do not provide statements of confidence about the importance of individual genes. Furthermore, using an arbitrary significance threshold to separate genes into adapted and non-adapted categories is obviously sensitive to the choice of an arbitrary parameter. If a gene in one species passed this threshold but fell just short in another, it would not contribute to the genome-wide signal of repeated adaptation. However, genome scans often result in continuous distributions of summary statistics that are not readily divisible into adapted/non-adapted categories. Such classification thus potentially screens out useful information in some lineages, particularly if genome scans are underpowered and/or when genes have weak signals of adaptation.

Here, we develop a method to estimate the involvement of an individual gene in repeated adaptation with less reliance on arbitrary thresholds. Provided we are considering adaptation to some specific selection pressure, we can make use of a simplifying assumption to compare evidence across multiple lineages: most genes in the genome are unlikely to be involved in the adaptative response. Under this assumption, a gene that is involved in adaptation will tend to fall into the tail of the genome-wide distribution, which can be represented by the rank of its summary statistic relative to the other genes in the genome. Comparisons of the rank-order of each ortholog of the gene across multiple lineages can then be used to assess the overall non-randomness of their summary statistics, which is therefore indicative of the gene’s involvement in repeated adaptation. Our method, which we call *PicMin*, uses the theory of order statistics to quantify the degree of non-randomness in measures of adaptation across multiple lineages for individual genes. *PicMin* is generalizable to any number of lineages and provides a *p*-value for each gene. We characterize the statistical power of *PicMin* and find that it can sensitively detect repeated adaptation even when genome scans have only weak power to detect adaptation in individual lineages, as long as multiple lineages are compared. We apply our new method to previously published genome scans performed on *Arabidopsis* species and find several genes that exhibit clear evidence for repeated adaptation. Note that throughout this paper, for simplicity we refer to “genes” as being the unit of analysis in genome scans, but in reality, genome scans are typically applied to non-coding portions of species’ genomes as well. Additionally, we use the term lineage rather than species throughout the rest of this paper as *PicMin* could be applied to distinct populations of the same species.

## Methods

### Identifying candidates for repeated adaptation using genome scan data

Our goal is to find genes or genomic regions that are used for adaptation to a similar environmental challenge in two or more lineages. Such patterns may suggest the repeated evolution of similar mechanisms in response to similar selective challenges. Our goal is distinct from asking whether a particular gene is used for adaptation in any of the lineages; if a gene were used by just a single lineage it may indicate its potential for evolutionary response to a particular selective challenge, but it would not indicate repeated adaptation. Thus, any test of repeated adaptation needs to incorporate the possibility that adaptation operates in just a single lineage into its null hypothesis. In comparisons of *n* lineages, we propose screening out genes that show no strong evidence of being used for adaptation in any of the lineages; there can be no repeated adaptation if there is no evidence of adaptation in any single lineage. For each of the remaining genes, we then remove the lineage with the strongest evidence of adaptation and test for repeated adaptation among any of the remaining *n* – 1 lineages. The null hypothesis for the test of repeated use is that none of the *n* –1 remaining lineages have used the gene for adaptation.

Let us assume that we have performed separate genome scans for adaptation in each of *n* lineages. These genome scans were performed such that we can obtain empirical *p*-values (*ep*-values) for tests of adaptation for *L* genes. An *ep*-value is the quantile of a given gene’s adaptation score relative to all other genes from that lineage. The *ep*-values reflect the strength of evidence against a null hypothesis of no adaptation, i.e., the *ep*-values for genes that are not involved in adaptation are drawn from a uniform distribution (*U*{0,1}). Each of the *L* genes has orthologs present in all lineages; we thus have a list of *n ep*-values for each gene. Because the procedure is intended to look for evidence of repeated adaptation, it is only relevant for cases where at least one lineage uses the gene for adaptation. The first step in our analysis is to remove genes that show no evidence for adaptation for at least one lineage. We do this by restricting our analysis to genes where at least one of the *n* lineages has an *ep*-value less than a threshold (*ɑ*_*Adapt*_).

If a gene passes this first screen, we ask “is there evidence that the distribution of *n –* 1 *ep*-values for the orthologs of the gene in the other lineages is shifted towards small values?” We test for a downward shift in the distribution of *ep*-values in the remaining *n* –1 lineages for individual genes using the theory of order statistics. By ordering the remaining *n* –1 *ep*-values from smallest to largest we obtain a set of order statistics. Under the null hypothesis that the *n* – 1 *ep*-values were generated from true null hypotheses, we have a set of *n* – 1 order statistics sampled from a uniform distribution. Following convention, order statistics are denoted *x*_(1)_, *x*_(2),_ … *x*_(n − 1),_ where *x*_(1)_ refers to the smallest (and first) value. Order statistics sampled from a uniform distribution have marginal distributions that belong to the beta distribution (Gentle 2009). These beta distributions have parameters *k* and (*ν* +1–*k*), where *ν* is the number of items in the list. In our case, there are *n* – 1 items in the list, so the marginal distributions have parameters *k* and *n-k*. Therefore the *k*^th^ order statistic from our set of *n* –1 *p*-values follows

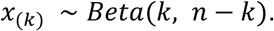

We can use the marginal distributions for each of the *n* – 1 order statistics to compute one-sided *p*-values as

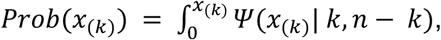

where *Ψ* (*k, n − k*) represents the probability density function of the beta distribution with parameters *k* and (*n*–*k*). We thus obtain ***X***, a list of *n* – 1 *p*-values (i.e. ***X* =** {***prob***(***x***_(***1***)_), ***prob***(***x***_(***2***)_), …, ***prob***(***x***_(***n***−***1***)_)}). The values in ***X*** represent individual hypothesis tests asking whether particular order statistics are smaller than expected by chance.

Multiple comparisons correction is then applied to the minimum of ***X*** to obtain a single *p*-value that reflects the evidence that a particular gene exhibits repeated adaptation. We use a multiple comparisons correction based on the method developed separately by Tippett (1931), Dunn (1958) and Šidák (1967). These methods assume independence among the tests being compared; however, the *p*-values obtained from the marginal distributions of order statistics are highly correlated. We therefore use the Tippett method as implemented in the *poolr* package for R (v1.1-1; Cinar and Viechtbauer 2022) to account for the dependency structure among the values in ***X***. We use the “empirical” Tippett method in *poolr*, which samples sets of *p*-values using an estimate of their dependency structure. To obtain the expected correlation matrix among values in ***X***, we simulate a set of *n p*-values drawn from a uniform distribution conditioning on at least one of them being smaller than *ɑ*_*Adapt*_ and replicate this procedure 10,000 times. We compute ***X*** for each of these simulated cases, then calculate the correlation matrix among values in ***X***. *poolr* samples *p*-values from such correlation matrices to build a null distribution against which to compare observed data. The rank of the observed data in this null distribution provides the combined *p*-value for a given gene. By default, we perform 100,000 replicates to build the null distribution, but more precise *p*-values can be obtained by performing more replicates.

A combined *p*-value obtained from this procedure can be used to test whether a particular distribution of *n* –1 order statistics is unusual under the null model. If that combined *p*-value were smaller than expected by chance (at a user-defined significance threshold that we call *ɑ*_*Repeated*_) it would provide evidence for a downward shift in the distribution of *p*-values and, since we are conditioning on there being evidence for adaptation in at least one of *n* lineages, evidence for repeated use of that gene for adaptation. From this procedure we also retain the index of the minimum value in ***X*** as it indicates the number of lineages that exhibit a pattern of repeated adaptation. For example, if we were applying *PicMin* to data from 7 lineages, ***X*** would contain 6 values for a gene. If, for a particular gene, the combined *p*-value was smaller than *ɑ*_*Repeated*_ and the minimum value in ***X*** was the 5th one, it would provide evidence for repeated adaptation at that locus for 6 out of the 7 lineages.

Real datasets will not necessarily include data for all genes. When analyzing empirical data with *PicMin* the user would need to obtain separate correlation matrices for each case represented in their data. For example, if a particular gene were only present in 13 out of 30 lineages, one would need to build a correlation matrix for a 13 lineage comparison to analyze that particular gene.

### Simulations

To characterize the performance of *PicMin* to detect repeated adaptation at a given locus, we simulate genome scan results for sets of 7 (or 30) lineages at individual genes. We simulated *ep*-values under the null hypothesis of no adaptation in any lineage by simply sampling ranks uniformly from integers up to 10,000 and dividing by 10,000, which represents a comparison of 10,000 genes across lineages. Genome scans will vary in their power to detect adaptation for numerous reasons (study design, population history, strength of selection, etc.); we use the term “genome scan power” to refer to the probability that a false null hypothesis results in an empirical *p*-value less than 5%. We simulated false null hypotheses for single genes and varied the genome scan power by sampling ranks from the integers from 1 to 500/*power* and dividing by 10,000. For example, to simulate false nulls in a genome scan with 50% power, we sampled ranks out of 1,000 and divided them by 10,000. In our simulations, we varied the total number of lineages being compared, the number of lineages exhibiting a false null and the genome scan power. We applied *PicMin* to identify repeated adaptation to the e*p*-values for each of these simulated datasets. In addition, we applied a binomial test on the number of lineages that passed *ɑ*_*Adapt*_, comparing the number of species that reject the null hypothesis for that gene to the expected number based on the Type I error rate. Due to computational limitations, we use a different approach to model genome-wide analyses.

To test the performance of *PicMin* to identify repeated adaptation in whole genome analyses, we simulated datasets of 10,000 genes across 7 lineages. In these simulations, genes that conformed to the null hypothesis of no adaptation had parametric *p*-values drawn from a uniform distribution (i.e., *U*{0,1}) while genes that exhibited adaptation had parametric *p-* values drawn from a leptokurtic beta distribution, using the “*rbeta(100, a, b)”* function in R. We chose the specific parameters of the beta distribution so that random draws would fall below 0.05 with a given probability. Table S1 lists the exact parameters we chose for this beta distribution and the corresponding genome scan power. We simulated repeated adaptation among 3, 5 or 7 out of 7 lineages for 100 genes, while the remaining 9,900 conformed to the null hypothesis. The parametric *p*-values for all 10,000 genes were converted into *ep*-values within each lineage. We applied *PicMin* to identify repeated adaptation to each gene in these datasets then applied a genome-wide false discovery rate correction using the *“p*.*adjust(*…, *method = “fdr”)*” command in *R*.

Note that we describe and evaluate *PicMin* assuming genome scans that result in *ep*-values, but if one had a suitable null model and was able to compute parametric *p*-values, those would be appropriate input to our method.

### Code availability

An implementation of our method to identify repeated evolution as well as detailed walkthrough documents are available at https://github.com/TBooker/PicMin. Code to perform all simulations, analyze data and generate all plots are also available at https://github.com/TBooker/PicMin.

## Results

### Choosing the best *ɑ*_*Adapt*_

Under the null hypothesis that only a single lineage used a particular gene for adaptation, the choice of *ɑ*_*Adapt*_ affects the shape of the distribution of *p*-values when we apply our method (Figures S1-2). When analyzing 7 lineages, setting *ɑ*_*Adapt*_ to a lenient value of 0.10 led to a deficit of false positives (i.e. a conservative test) while particularly stringent *ɑ*_*Adapt*_ values led to excess false positives (Figure S1,S3). Setting *ɑ*_*Adapt*_ at 0.05 provided a test that had a uniform distribution of *p*-values under the null hypothesis in this case (Figure S1). When analyzing 30 lineages, more stringent *ɑ*_*Adapt*_ values yielded uniform distributions of *p*-values under the null (Figure S2). The reason for this difference is that as the number of lineages being compared increases, the probability of at least one lineage passing *ɑ*_*Adapt*_ in a comparison increases. Removing the *ɑ*_*Adapt*_ screen altogether resulted in non-uniform distribution of *p*-values with a pronounced deficit of small *p*-values (Figure S1-2).

### Performance of *PicMin* to identify repeated adaptation

*PicMin* is a powerful test to identify candidates for repeated adaptation when the number of lineages exhibiting repeated adaptation is close to the total number of lineages tested. Under the null hypothesis that only a single lineage exhibits adaptation, *PicMin* gave the expected proportion of false positives (Figure 1). In a comparison of 7 lineages, *PicMin* had very high power to reject the null hypothesis when there were 5 or more lineages exhibiting a pattern of repeatability, even when the underlying genome scans had low statistical power (Figure 1A). In datasets comparing 30 lineages, we found that *PicMin* had almost perfect power to detect repeatability when the number of lineages exhibiting repeated adaptation is close to the total number of lineages being compared (Figure 1B). On the other hand, *PicMin* had very little power to detect repeated adaptation when the true number of lineages exhibiting repeated adaptation was small (Figure 1). For example, when comparing 7 lineages, there was virtually no power to detect repeated adaptation when only a pair of lineages exhibits repeated adaptation (Figure 1). *PicMin* had much higher power to detect repeated adaptation than a binomial test on the number of lineages with genes whose *ep-*values passed *ɑ*_*Adapt*_ (Figure S4). Statistical power to detect repeated adaptation with *PicMin* is largely insensitive to the choice of *ɑ*_*Adapt*_, though more stringent *ɑ*_*Adapt*_ values lead to a slight increase in power when around half of all lineages tested exhibit a pattern of repeated adaptation (Figure S5).

**Figure 1.**
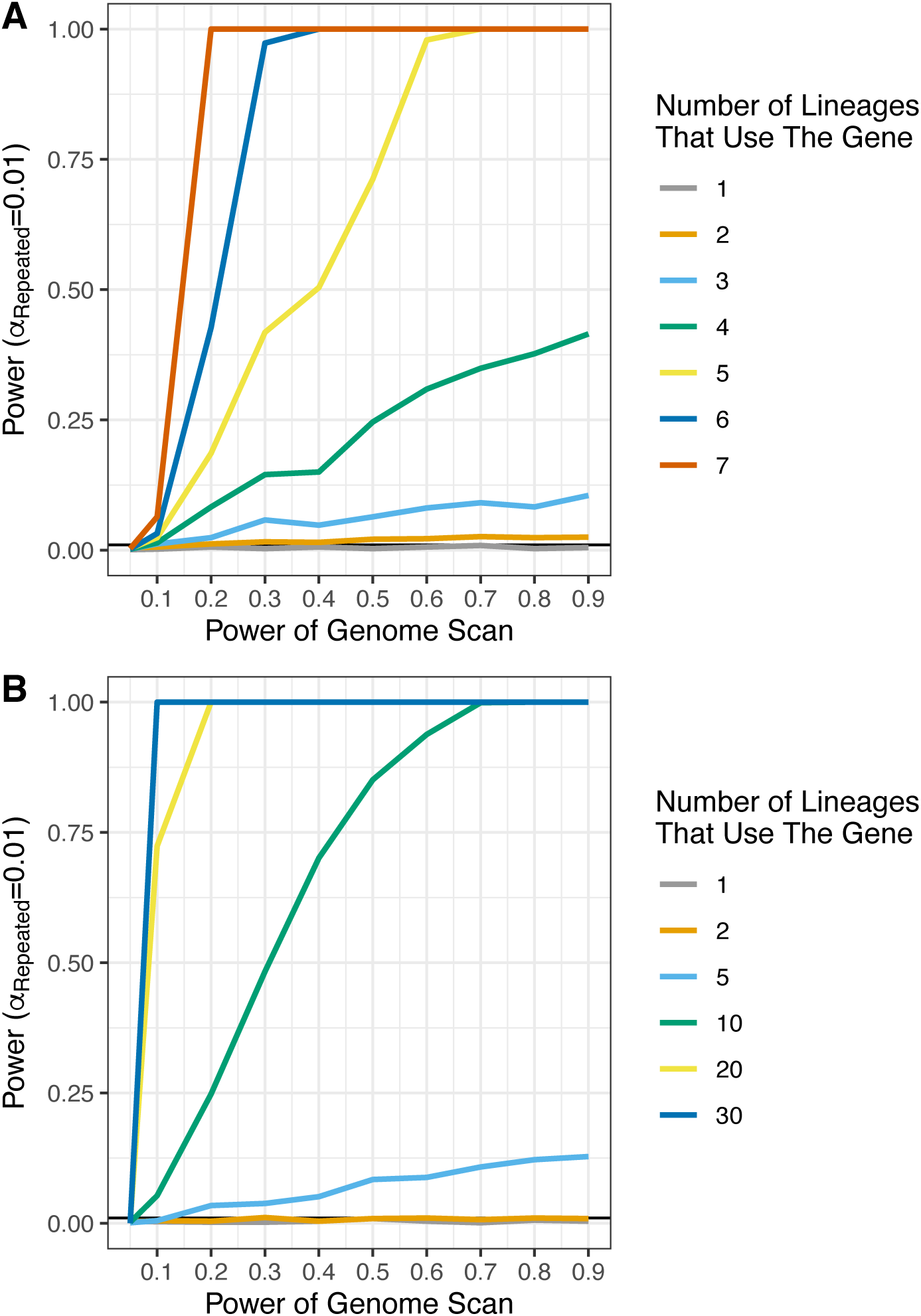
Statistical power of *PicMin* to identify repeated adaptation from multiple genome scan datasets. Panel A) shows results for a study containing 7 lineages and B) shows results for a study of 30 lineages. Simulations of individual genes were performed conditioning on there being at least one lineage with a *p*-value less than *ɑ*_*Adapt*_. For comparisons of 7 lineages *ɑ*_*Adapt*_ = 0.05, but for 30 lineages *ɑ*_*Adapt*_ = 0.01. The expected false positive rate (*ɑ*_*Repeated*_) is indicated as a solid horizontal black line. For visualization purposes, we only show the results for a subset of all possible configurations in panel B.

### How many lineages exhibit repeated adaptation?

*PicMin* provides an estimate of the number of lineages that exhibit a pattern of repeated adaptation. In Figure 2, we compare the estimates with our new method to those obtained by simply counting the number of lineages passing *ɑ*_*Adapt*_. The results shown in Figure 2 correspond to different combinations of genome scan power and numbers of lineages exhibiting repeated adaptation. When only 3 lineages out of 7 used a particular gene for adaptation and genome scans had low power, estimates of the number of lineages exhibiting repeated adaptation were particularly inaccurate. When 5 or 7 lineages used the gene for adaptation, estimates of the number of lineages exhibiting repeatability from our new method were much more accurate than using a simple fixed threshold scheme. This difference was particularly striking when genome scans had weak power in individual lineages (left columns in Figure 2).

**Figure 2.**
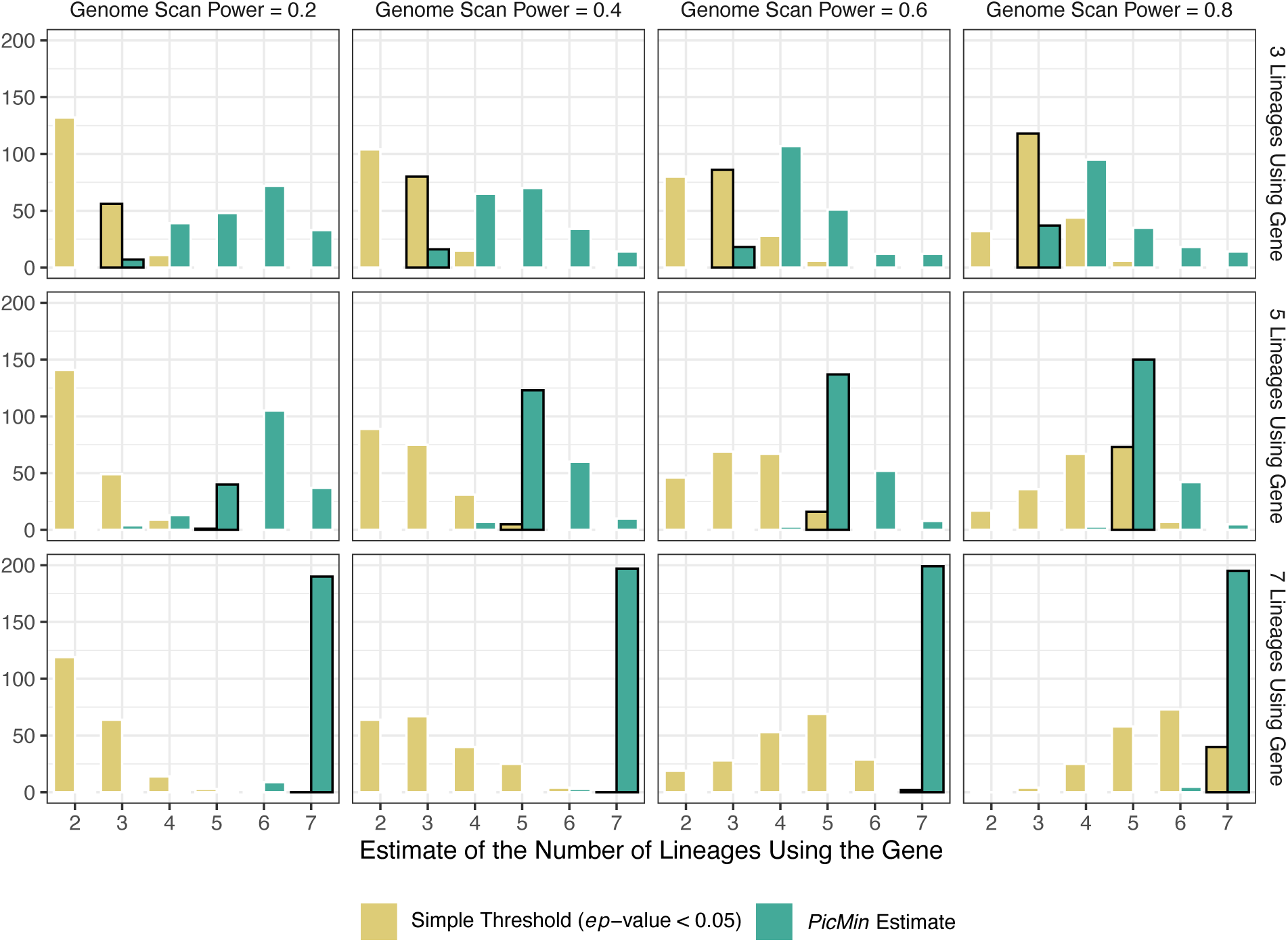
Estimates of the number of lineages exhibiting a pattern of convergence. The true number of lineages using the gene is highlighted with a black outline. The graphs shows results only from cases that gave significant (*p* < 0.05) evidence of repeated adaptation. Each panel in the plot shows the results from 200 simulation replicates. The “simple threshold” shows the number of lineages with a *ep*-value < 0.05 for the gene.

### Performance of *PicMin* to identify repeated evolution in genome-wide analyses

When attempting to identify regions of the genome that exhibit repeated adaptation, researchers will likely perform many simultaneous hypothesis tests on many genes or genomic windows. These multiple comparisons require adjusting for the number of hypotheses tested, in order to have confidence in the result for any given gene. We performed simulations to test how well *PicMin* performs in the context of whole genome analysis of 7 lineages. In these simulations, we modeled genome scans that had been performed on 10,000 genes present in all 7 lineages, where there were 100 true positives. Figure 3 shows the total number of significant genes that were identified after false discovery rate correction in cases with varying genome scan power and numbers of lineages exhibiting repeated adaptation. False discovery rate correction was applied using *p*.*adjust(*…, *method = “fdr”)* in R, which implements the Benjamini-Hochberg method (Benjamini and Hochberg 1995). The method described here was able to identify repeated adaptation among 7 out of 7 lineages even when individual genome scans had weak power to detect adaptation in individual lineages (Figure 3). When only 5 out of 7 lineages exhibited repeated adaptation, *PicMin* was able to identify true positives when genome scans in individual lineages had high statistical power (Figure 3). When only 3 out of 7 lineages exhibited repeated adaptation, *PicMin* was not able to identify true positives at genome-wide significance (Figure 3). Across all parameter combinations, the number of false positives was very low (Figure 3).

**Figure 3.**
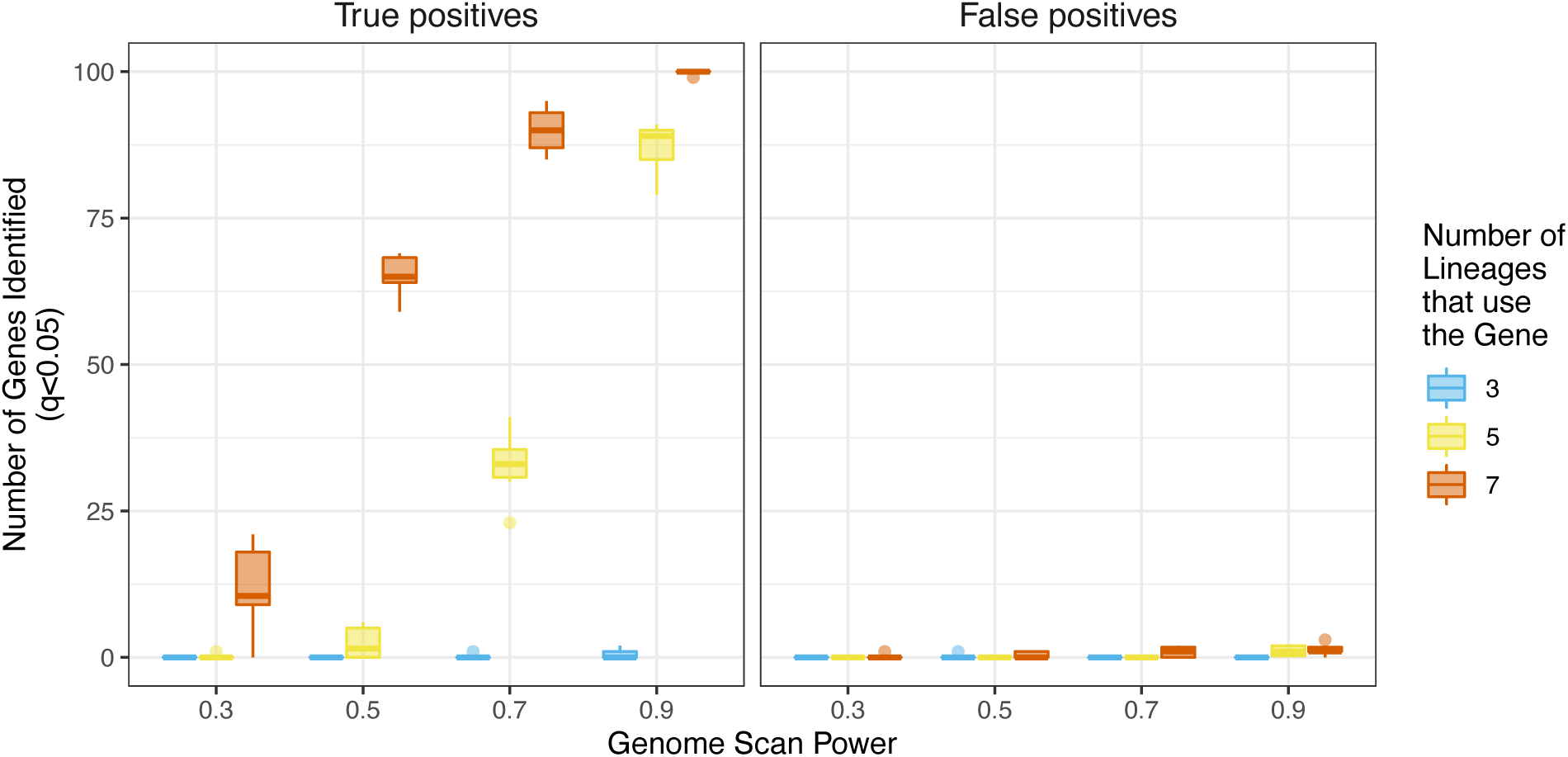
The total number of genes with evidence for repeated adaptation identified in a comparison of 7 lineages after applying a genome-wide false discovery rate correction (*q* < 0.05). Boxes show the results from 10 replicate analyses. Panel A shows the total number of true positives identified and B shows the total number of false positives in each case. We set *ɑ*_*Adapt*_ = 0.05 for these simulations.

### Application of *PicMin* to genome scans in *Arabidopsis spp*

As an empirical test of *PicMin*, we re-analyzed results of genome scans in *Arabidopsis* species originally reported by Bohutínská et al. (2021). Bohutínská et al. (2021) generated population genomic data for lineages that have each independently colonized alpine environments (2 from *A. halleri* and 5 from *A. arenosa*). For each lineage, Bohutínská et al. (2021) calculated *F*_*ST*_ between foothill and alpine population pairs in non-overlapping 1 Kbp analysis windows across the genome. We performed *PicMin* on all windows that had data for at least 4 lineages and applied a genome-wide false discovery rate correction to the resulting *p*-values. We identified 10 loci that had significant evidence genome-wide (*q* < 0.05) for repeated adaptation, of which 3 were in tight linkage (Figure S6). These loci overlapped 5 genes, 3 of which had been identified previously by Bohutínská et al. (2021). For two of the genes identified, the estimated number of lineages exhibiting repeated adaptation from *PicMin* was larger than had been estimated by either a fixed threshold-based approach or the approach used by Bohutínská et al. (2021)(Table S2). Using a restrictive approach (*q* < 0.05) we identified 5 genes with evidence for repeated adaptation genome-wide, whereas Bohutínská et al. (2021) used a more inclusive approach and identified 151 genes. If we adopt a more inclusive approach, using a significance threshold of *q* < 0.5, 346 loci were identified by *PicMin* corresponding to 167 genes. Of those genes, 101 were also identified by Bohutínská et al. (2021). For those genes, the average number of lineages implicated in repeated adaptation estimated by *PicMin* was 5.23 but only 2.04 using the approach of Bohutínská et al. (2021). The mean number of lineages implicated in repeated adaptation obtained by simply counting the number of lineages with *F*_*ST*_ values in the 95^th^ percentile (i.e. *ep-*values < 0.05) of each lineages respective distributions for these genes was 2.99.

## Discussion

In this paper, we have described a novel method to identify regions of the genome that exhibit patterns consistent with repeated adaptation. If one observes repeated adaptation at a particular gene, it not only sheds light on interesting aspects of evolution, but it serves to confirm the results of genome scans in individual species, which have notorious problems with disentangling how the interaction between space, drift, and selection affects false positives and negatives (Hoban et al 2016; DeRaad et al. 2020). Because drift is unlikely to drive a repeated pattern of association to environment across multiple species, *PicMin* can provide stronger evidence about adaptation than is possible from individual genome scans (provided other factors that may drive repeated associations are controlled, as discussed below). In the Introduction, we outlined how a statistical test for repeated evolution would ideally make use of continuous variation rather than relying on arbitrary thresholds. Our new method screens out genes that are unlikely to be contributing to adaptation, and within the remaining set uses the quantitative evidence for adaptation in the other lineages to obtain a statement of evidence for repeatability. *PicMin* is able to detect repeated adaptation at a locus even when genome scans in individual lineages have low power (Figure 1, Figure 3). Estimates of the number of lineages exhibiting a pattern of repeated adaptation obtained from *PicMin* are often more accurate than those made using a simple threshold-based approach (Figure 2).

Statistical tests of repeated adaptation should account for the possibility that just a single lineage exhibits a strong signal. While we were developing *PicMin*, a very similar, but general purpose meta-analysis method based on order statistics was published by Yoon et al. (2021). Their method, *ordmeta* (Yoon et al. 2021), asks whether the combined evidence from all the tests is sufficient to reject a common null hypothesis. Such an approach would allow one to ask whether any of a set of lineages used a gene to contribute to adaptation. However, this is a different question from asking whether a gene is repeatedly used by more than one lineage, the target of our current approach. The question of repeated adaptation is addressed by (1) removing genes that are unlikely to be contributing to adaptation in any lineage, and (2) after dropping the lineage with the strongest evidence of adaptation, using a modified order statistics approach on the remaining lineages (accounting for the change in the correlation structure of the list of *p-*values caused by that dropping of one non-random lineage). If one were to use *ordmeta*, or other combined probability approaches, to detect repeated adaptation without accounting for the null hypothesis of just a single lineage exhibiting adaptation, the false positive rate becomes higher than stated.

Other methods to study repeated adaptation have focused on the genome scale, comparing the observed number of genes that have signatures of adaptation in multiple lineages with the expectation for a random draw from the genes with signatures of adaptation in each lineage, as implemented in the *SuperExactTest* (Wang et al. 2015) and the C-score statistic (Yeaman et al. 2018). Such approaches yield a set of genes that are all potentially involved in repeated adaptation, but do not assess the statistical significance of individual genes. By contrast, the *PicMin* method developed here assigns statistical significance to individual genes, and is powerful when the number of lineages and underlying repeatability is sufficiently large. Depending on the application, *PicMin* could be deployed as a high-stringency test to find the genes with strongest evidence of repeated adaptation (by setting a stringent false discovery rate cutoff) or as a permissive screen to identify a set of genes that can then be studied as a group to test for enrichment of other features, such as gene ontology terms or differential expression. In the example here using the dataset of Bohutínská et al. (2021), we identified 5 genes at a restrictive *q* < 0.05 (Table S2) or 167 genes at a more inclusive *q* < 0.5.

Despite low power to identify individual genes in an analysis of a pair of lineages (Figure 1), *PicMin* can still be used to assess genome-wide evidence for repeated adaptation. By relaxing the stringency used when identifying candidates for repeated adaptation, one could use *PicMin* to obtain an estimate of the number of lineages exhibiting repeated adaptation. For example, if one had *ep*-values corresponding to *L* gene orthologs for a pair of lineages, the first step would be to screen out genes where neither of the lineages has an *ep*-value ≤ *ɑ*_*Adapt*_. The number of genes that pass this initial screen where the larger of the two *ep*-values ≤ *ɑ*_*Repeated*_ provides an estimate for the number of genes that have evidence for repeated adaptation. The number of false positives expected from such a 2-way analysis is easily calculated as 2***a***_-./01_***a***_23%45675_*L*. This procedure could be applied to specific pairs of lineages; for example, to test the hypothesis that the extent of repeatability decays with phylogenetic distance similar to the analysis of Bohutínská et al. (2021).

### The value of empirical *p*-values

Consider the problem of determining the genetic basis of local adaptation. Genome scans commonly assess evidence for local adaptation based on the strength of correlation between allele frequency and environmental variables across multiple populations; genotype-environment association (GEA) analyses. Most species live in spatially structured environments where genetic drift and restricted migration drives a pattern of isolation by distance (Wright 1949). If environmental variables covary with a pattern of isolation by distance, many alleles will exhibit associations with environmental variation and simple GEA analyses may often result in false positives. There are multiple methods that build population structure into GEA analyses (reviewed in Hoban et al. 2016), but these can result in false negatives if the true drivers of adaptation have spatial patterns in allele frequency that align with population structure (DeRaad et al. 2021). However, under the assumption that most genes in the genome are not contributing to adaptation, the rank-order of GEA summary statistics (e.g. *p*-values or Bayes factors from correlation tests) should result in enrichment of the causal loci in the lower tail of the genome-wide distribution (Hancock et al. 2011). If bouts of local adaptation in other lineages are independent, it is unlikely that a non-causal gene will tend to fall into the tail of this distribution in multiple lineages, and so a test can be developed to assess evidence for a gene’s importance using these rank-order distributions. This approach of converting summary statistics into empirical *p*-values (or as we refer to them in this paper, *ep*-values) based on their rank-order has been used previously to represent the relative strength of evidence for a gene’s involvement in local adaptation (Hancock et al. 2011) and as the basis for identifying outliers from genome scans; percentile thresholds are often applied to the empirical distribution of summary statistics to identify candidates for adaptation.

### A strategy for handling paralogs

Identifying repeated adaptation requires an understanding of orthology and paralogy among the lineages being compared. The recent studies of repeated adaptation by Bohutínská et al. (2021) and Tittes et al. (2021) analyzed species or lineages that were closely enough related that a single reference genome was suitable for all samples. Using a single reference genome for all samples makes identifying loci that exhibit repeated adaptation fairly straightforward, but there are potential complications. In some lineages, ancestral genome duplications and/or rearrangements may have caused paralogs to be located in dramatically different genomic regions. If different paralog copies had signals of adaptation in different lineages, that may still be considered evidence for repeated adaptation. This issue of paralogy/orthology will be particularly pronounced when analyzing distantly related species where there is little synteny among genomes being compared. In such cases, one may identify gene orthologs present in different lineages using packages such as *OrthoFinder* (Emms and Kelly 2019). *OrthoFinder* attempts to allocate genes into orthogroups, groups of genes that were inherited from a single copy in the common ancestor of the lineages being analyzed. Comparing evidence for adaptation among orthogroups, rather than individual genes, may be a useful way to identify repeated adaptation.

A strategy for dealing with paralogs would be to combine the *ep*-values for the genes belonging to each orthogroup within a lineage, whilst correcting for multiple comparisons. For a particular lineage possessing *y* members in orthogroup *z*, one could obtain a combined *ep*-value using the Tippett-Dunn-Šidák correction:

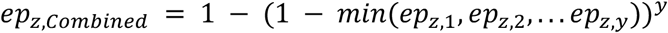

These combined *ep*-values for each orthogroup could then be compared across lineages using *PicMin* for identifying repeated adaptation. Combining information within orthogroups will decrease statistical power but given that the method we have developed can identify repeated evolution from weakly powered genome scans (Figure 1), such a method may be useful for comparing distantly related lineages.

### Repeated patterns without repeated adaptation

At root, the method we have developed is not actually testing a null hypothesis of no adaptation–there is no evolutionary model built into our method. Rather, *PicMin* is testing for a repeated pattern of a signal correlated with adaptation. Such a pattern could come from shared adaptation from de novo mutations, parallel evolution based on shared genetic variation, or shared genetic signatures of ancient selection that were inherited from a common ancestor, and not all of these may be the desired pattern of adaptation that is the target of the study. Furthermore, if the tested lineages experience ongoing hybridization, repeated signals of association could be driven by introgression. It is important to keep in mind that any factor that drives extreme test statistics in orthologous genomic regions across lineages could resemble the pattern expected by repeated adaptation. There are several processes of this kind that are worth mentioning. Firstly, genetic hitchhiking may cause linked genomic elements to exhibit similarly extreme summary statistics in genome scans. If there was conservation of synteny among the lineages being compared, true repeated adaptation at a single locus may cause a signal of adaptation at neutral linked genes in multiple lineages. We found a result consistent with this effect in the analysis of data from Bohutínská et al. (2021), where 3 contiguous windows and 1 closely linked window all had significant evidence for repeated adaptation (Figure S6; Table S2). The gene overlapping or closely linked to these outliers has previously been reported to harbor trans-specific polymorphisms in *Arabidopsis spp*. (Guggisberg et al. 2018). Furthermore, background selection is likely ubiquitous across eukaryotic genomes, with regions of the genome with high functional density and low recombination rates potentially subject to extreme background selection effects (reviewed in Comeron 2017). If the genome scan methods used are sensitive to background selection, shared profiles of functional density and recombination rate could lead to false signals of apparent adaptation repeatedly across lineages (Burri et al. 2015). In species that exhibit wide variation in recombination rates across the genome, outliers in scans for adaptation may tend to occur in genomic regions with low recombination rate (Booker et al. 2020). If recombination rate landscapes are conserved across the lineages being compared, outliers may occur in similar genomic regions due to the effects of low recombination yielding patterns that resemble repeated adaptation (Booker et al. 2020). As with most genome scans, searching for repeated adaptation using the methods we outline in this paper should be treated as the first step in a process of hypothesis generation, and alternative sources of information must be used to confirm that these signals are indeed caused by selection and adaptation.

## Acknowledgements

Thanks to Magdalena Bohutínská for access to and help with the *Arabidopsis* genome scan results as well as comments on the manuscript. Thanks to Pooja Singh, James Whiting and attendees of the delta-tea discussion group at UBC for helpful discussions. TRB is supported by a Bioinformatics Postdoctoral Fellowship awarded by the Biodiversity Research Centre at UBC. Funding for this work was provided by Genome Canada, Genome Alberta and NSERC Discovery Grants awarded to MCW and SY. SY is supported by an AIHS research grant. This study is part of the CoAdapTree project which is funded by Genome Canada (241REF), Genome BC and 16 other sponsors (http://coadaptree.forestry.ubc.ca/sponsors/).

## Supplementary Material

**Table S1.** Parameters of the beta distribution used to simulate *p*-values under adaptation for simulations of whole genome datasets.

| Parameter of the beta distribution |  | GEA Power |
| --- | --- | --- |
| $a$ | $b$ | |
| 0.8 | 5 | 0.3 |
| 0.5 | 5 | 0.5 |
| 0.3 | 5 | 0.7 |
| 0.1 | 5 | 0.9 |

**Table S2.**
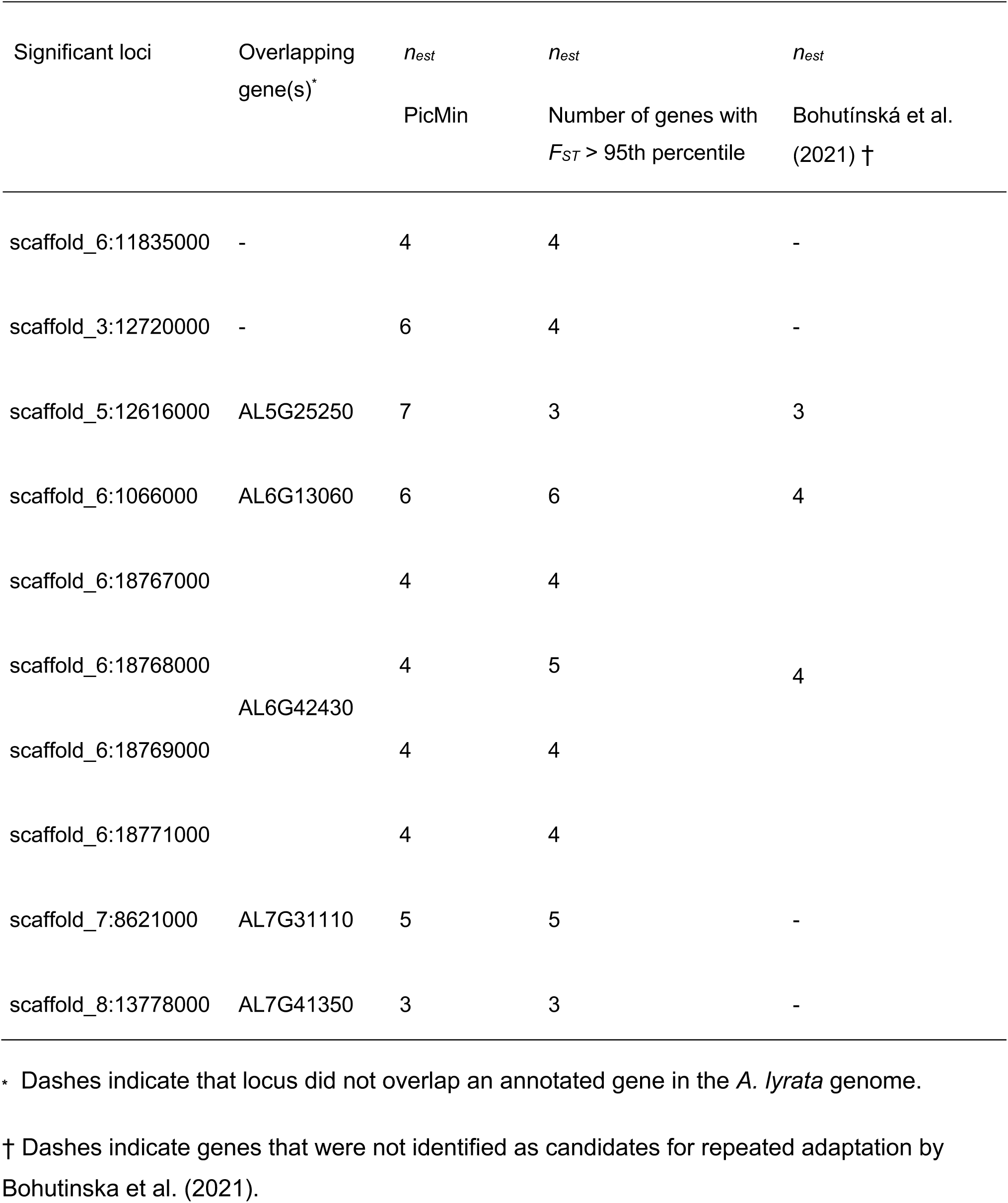
Analysis windows identified as having evidence for repeated adaptation (*q*<0.05) from the *F*_*ST*_ results of Bohutínská et al. (2021). Columns labelled *n*_*est*_ indicate the estimate of the number of lineages implicated in repeated adaptation from three different tests. Results for *n*_*est*_ from et al. (2021) were obtained from their Supplementary Dataset S4.

**Figure S1.**
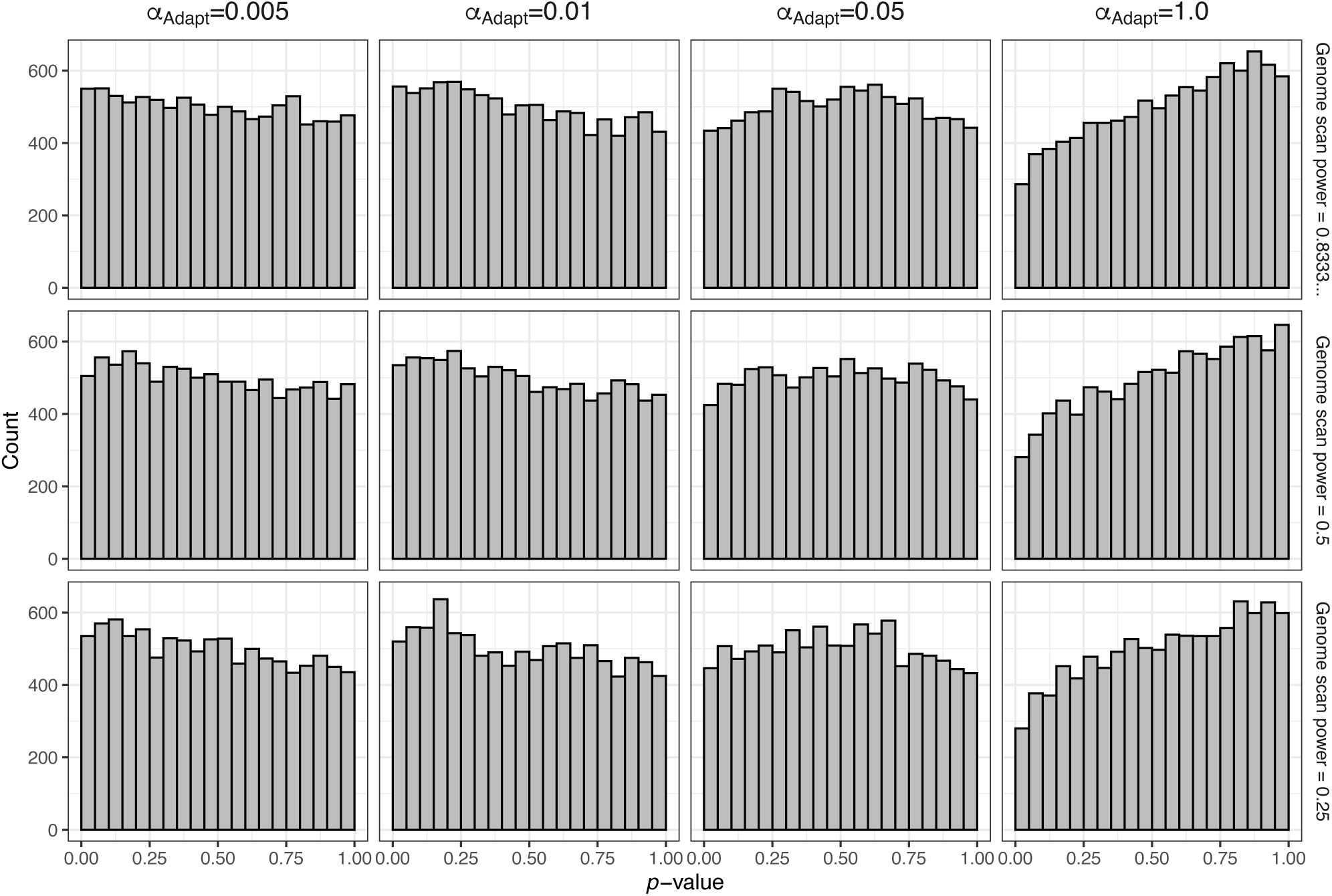
The distribution of *p*-values from *PicMin* under the null hypothesis that only one species uses a particular gene for adaptation. Results for comparisons of 7 lineages are shown. 10,000 replicates analyses are shown in each panel.

**Figure S2.**
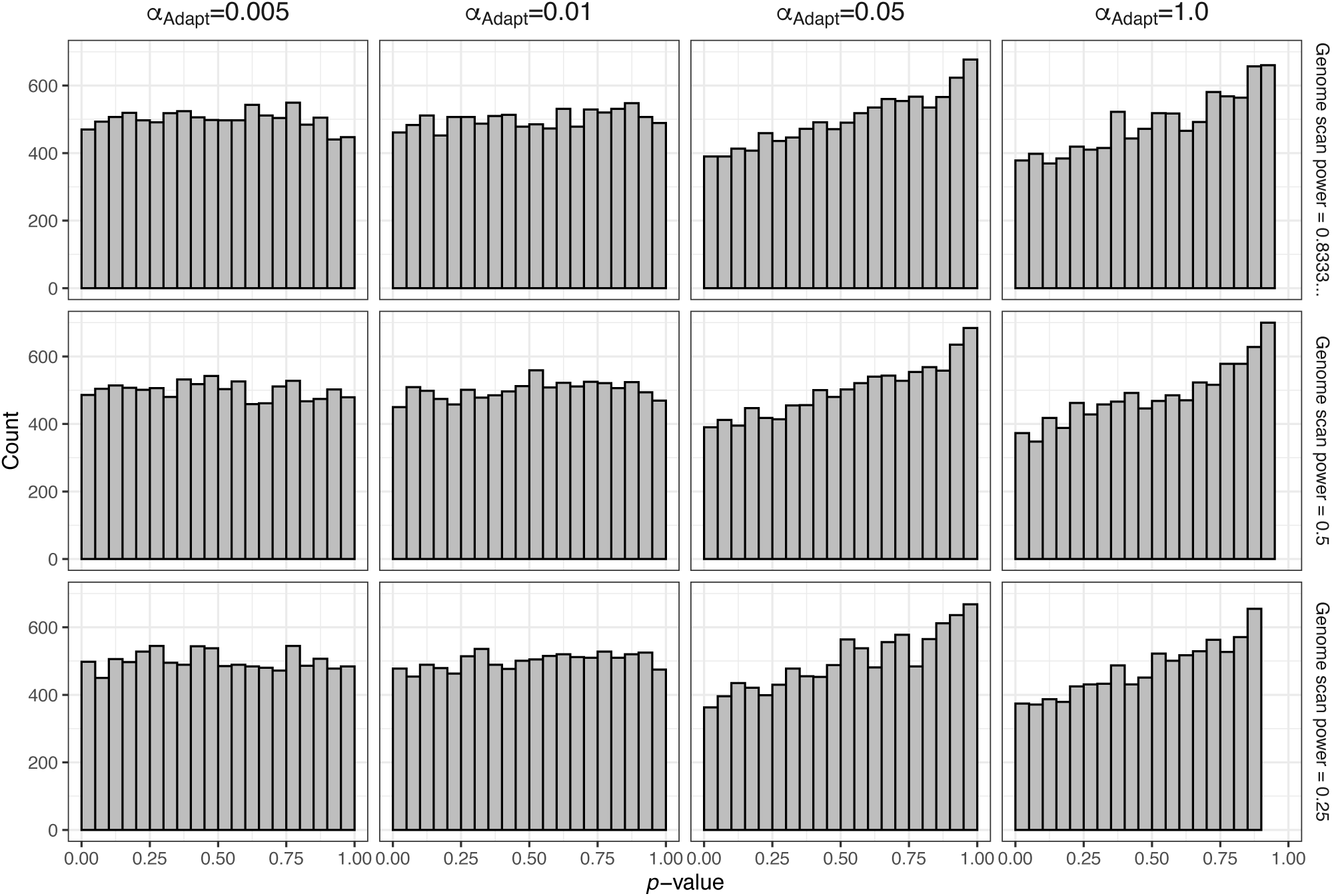
The distribution of *p*-values from *PicMin* under the null hypothesis that only one species uses a particular gene for adaptation. Results for comparisons of 30 lineages are shown. 10,000 replicates analyses are shown in each panel.

**Figure S3.**
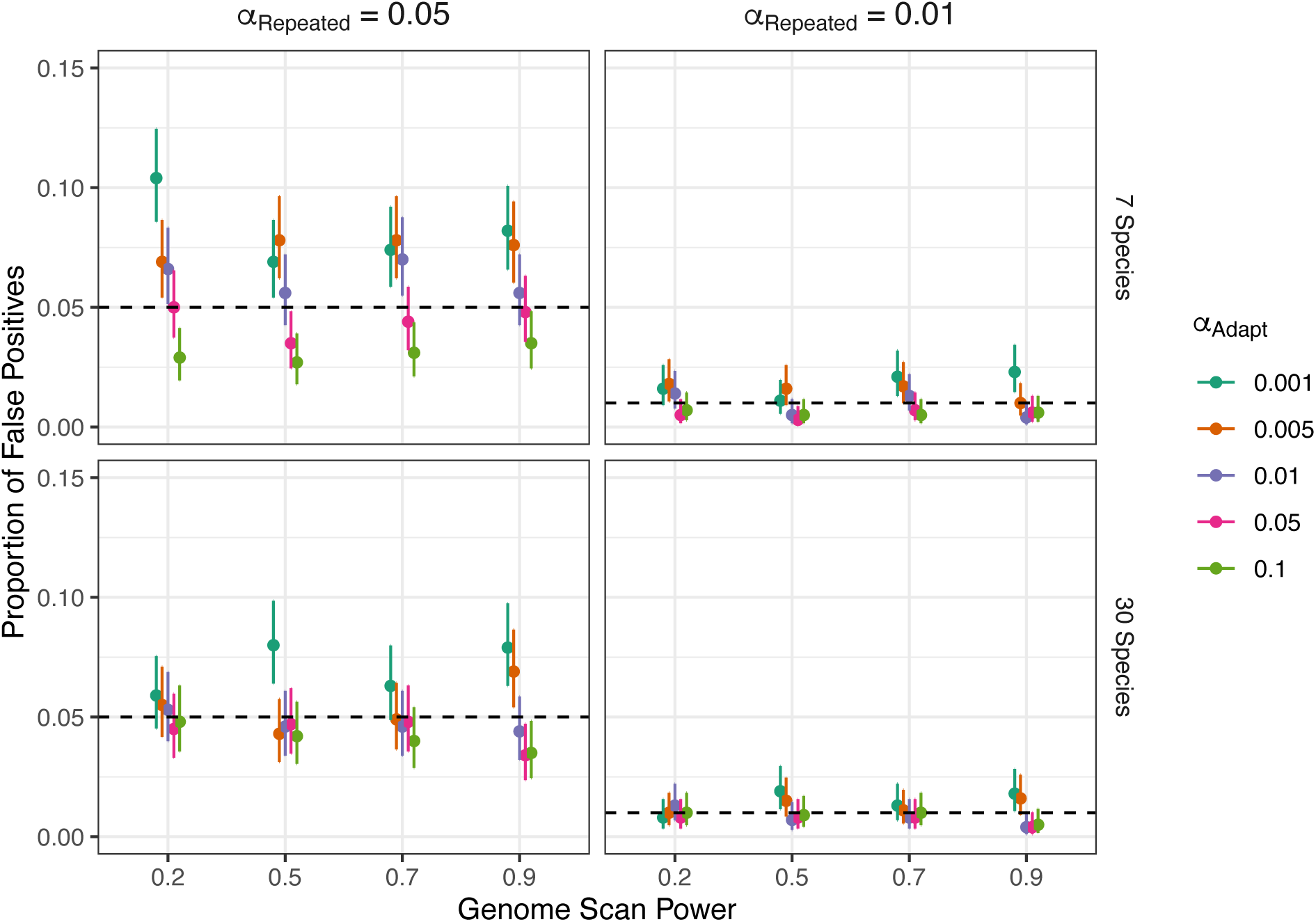
The choice of *ɑ*_*Adapt*_ on false positive rates. Error bars represent 95% Clopper-Pearson intervals.

**Figure S4.**
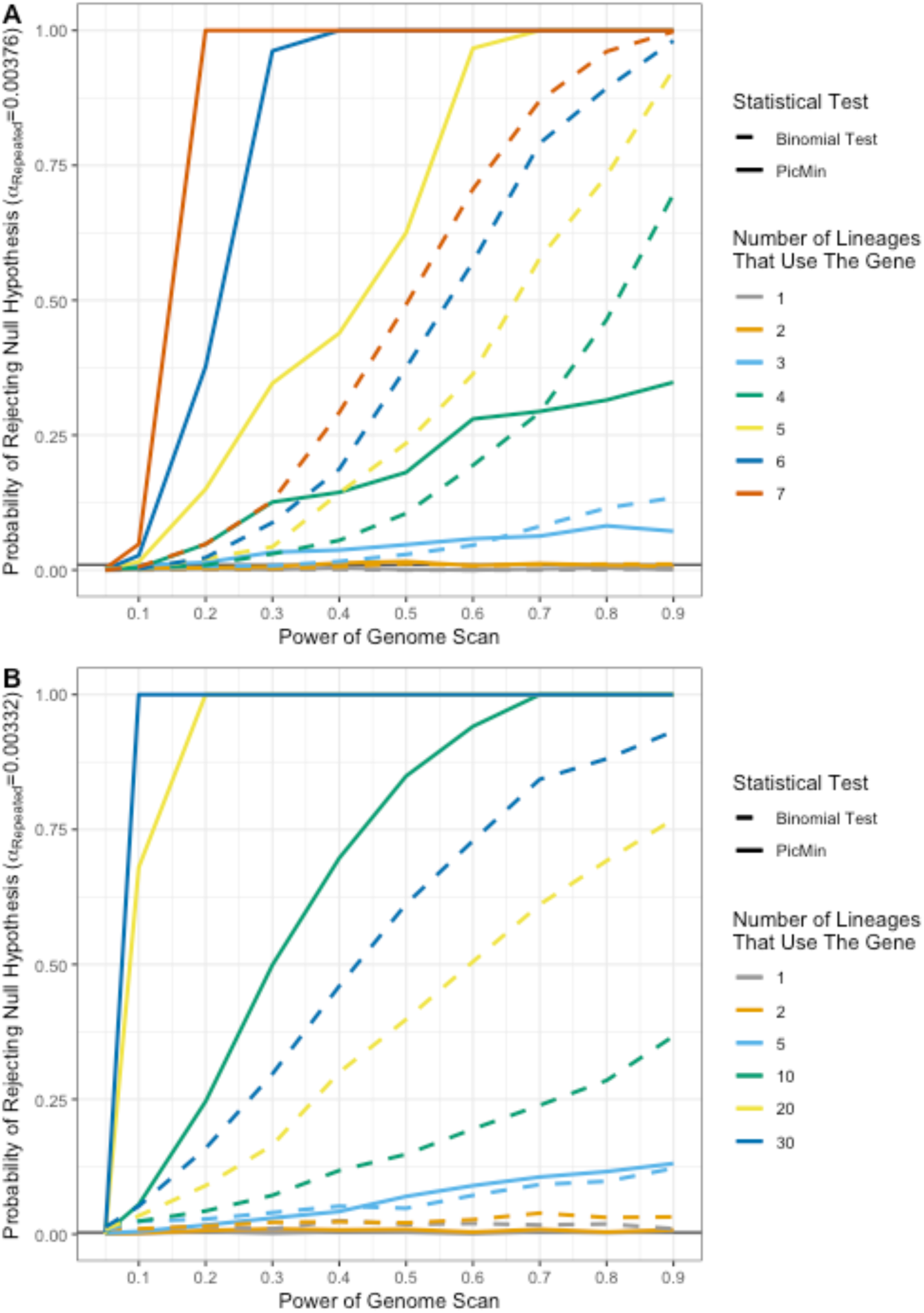
Comparison of *PicMin* and the binomial test to identify repeated adaptation from multiple genome scan datasets. Panel A) shows results for a study containing 7 species and B) shows results for a study of 30 species. Simulations of individual genes were performed conditioning on there being at least one lineage with a *p*-value less than *ɑ*_*Adapt*_ = 0.05. The expected false positive rate (*ɑ*_*Repeated*_) is indicated as a solid horizontal black line. For visualization purposes, we only show the results for a subset of all possible configurations of repeated adaptation in panel B. The binomial test computes the probability that *k* or more lineages (out of *n* lineages total) would have *ep* < *ɑ*_*Adapt*_ under a random draw, using “*binom*.*test(k, n, ɑ*_*Adapt*_)” in R.

**Figure S5.**
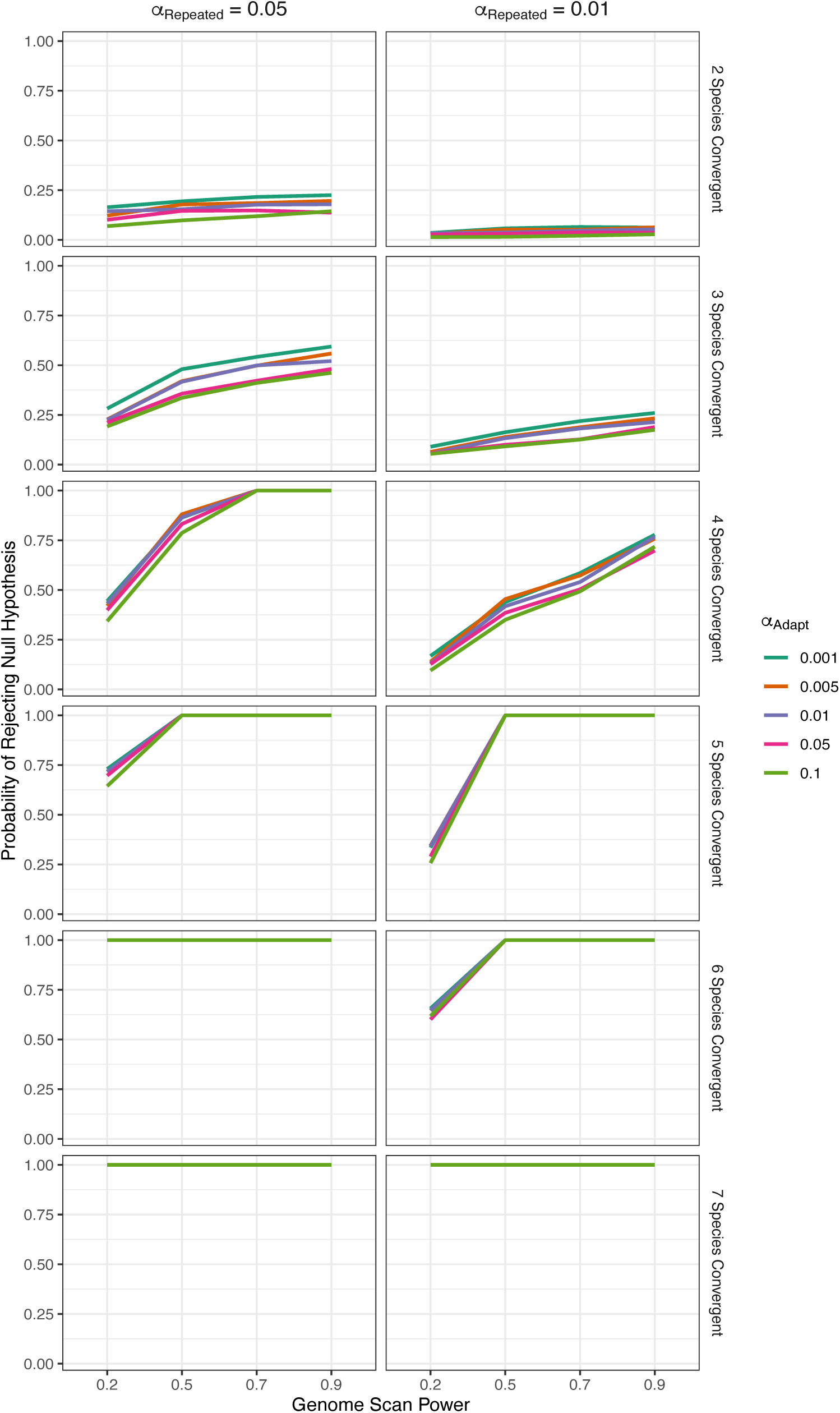
The effect of the choice of *ɑ*_*Adapt*_ on statistical power. Results are shown for a 7 species comparison.

**Figure S6.**
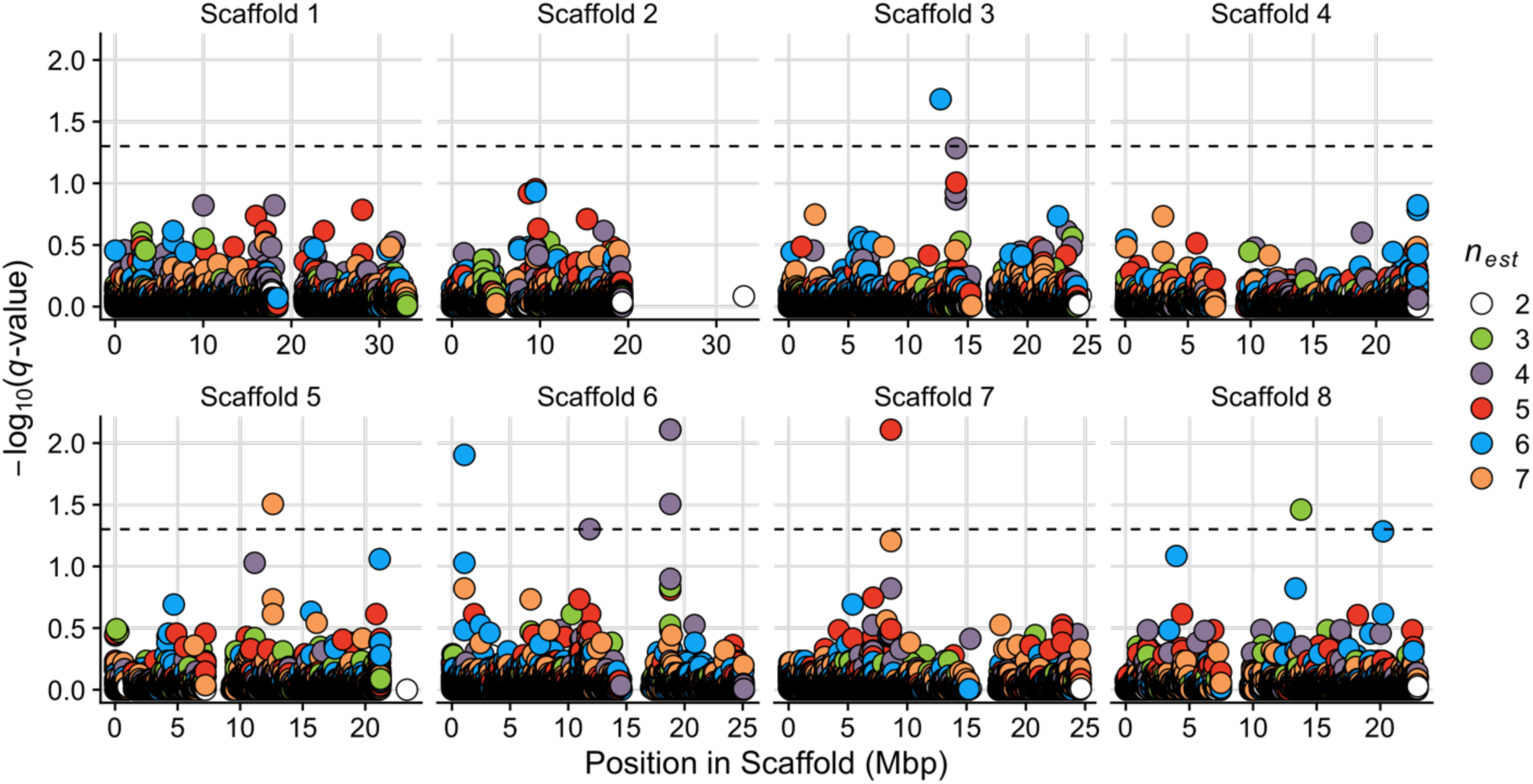
The results of *PicMin* as applied to the comparison of 7 species/populations of *Arabidopsis arenosa* and *A. halleri* reported by Bohutínská et al. (2021). We set *ɑ*_*Adapt*_ = 0.05 for these simulations. The dashed line indicates a genome-wide significance threshold of *q =* 0.05. *n*_*est*_ refers to the estimated number of lineages exhibiting a pattern of repeated adaptation.

